# Adolescent rats extend help to outgroup members, highlighting a neural network for group identity categorization

**DOI:** 10.1101/2021.11.29.470434

**Authors:** Jocelyn M. Breton, Jordan S. Eisner, Vaidehi S. Gandhi, Natalie Musick, Aileen Zhang, Kimberly L.P. Long, Olga S. Perloff, Kelsey Y. Hu, Chau M. Pham, Pooja Lalchandani, Matthew K. Barraza, Ben Kantor, Daniela Kaufer, Inbal Ben-Ami Bartal

**Author notes:** Department of Psychiatry, Columbia University, New York, NY, 10027, USA. Department of Psychiatry and Behavioral Sciences, University of California San Francisco, San Francisco, CA, 94143, USA.

## Abstract

Prosocial behavior, in particular helping others in need, occurs preferentially in response to the perceived distress of one’s own group members, or ingroup. The development of neural mechanisms underlying social selectivity towards ingroup members are not well established. Here, we used a rat helping behavior test to explore the development and neural basis of ingroup bias for prosocial behavior in adolescent rats. We previously found that adult rats selectively help others from their own social group, and that this selectivity is associated with activation in reward and motivation circuits. Surprisingly, we found that adolescent rats helped both ingroup and outgroup members, evidence suggesting that ingroup bias emerges in adulthood. Analysis of brain-wide neural activity, indexed by expression of the early-immediate gene c-Fos, revealed increased activity for ingroup members across a broad set of regions, which was congruent for adults and adolescents. However, adolescents showed reduced hippocampal and insular activity, and increased orbitofrontal cortex activity compared to adults. Adolescent rats who did not help trapped others also demonstrated increased amygdala connectivity. Together, these findings demonstrate that biases for group-dependent prosocial behavior develop with age in rats and suggest that specific brain regions contribute to this prosocial selectivity, overall pointing to possible targets for the functional modulation of ingroup bias.

**One Sentence Summary:** Prosocial selectivity increases with age in parallel with hippocampal and insular activation, providing insight into the neural classification of group membership.

## Introduction

Responding to another’s distress with a prosocial action is a crucial component of life in social groups[1-3]. Distress is a salient signal that can elicit empathy in the observer and recruit motivational responses intended on helping the distressed individual[4]. Yet, the empathic response to distressed others is largely impacted by their social identity, and prosocial behavior, in humans as well as other species, is selective and preferentially extended to affiliated others [5, 6]. To put it simply, we are much more likely to help others we care about than those we do not. For humans and other social species, affiliation expands beyond individual familiarity to encompass others of the same social group, or ingroup members[7, 8]. Identifying others as ingroup or outgroup members thus comprises a quick, effective heuristic for determining prosocial motivation. This mechanism, ostensibly adaptive at its origin, is hugely detrimental in modern society. Yet social bias, in particular with regard to prosocial motivation, is difficult to influence[9]. Targeting the formation of social bias during development, when social information is especially salient[10, 11] yet flexible[12], is a promising strategy for influencing behavior towards outgroup members along the life span, and understanding the neural mechanisms underlying the development of social bias, is thus a key question of our time.

Evidence suggests that children categorize others into ingroup and outgroup members and demonstrate social preferences very early in development [13, 14]. For instance, babies prefer faces of same-race adults [15] or adults with the same accent as their caretakers [16], and use group membership information to guide behavioral choices [17, 18]. By 3-4 years of age, children can show ingroup favoritism [19, 20], even towards arbitrarily determined groups [21]. However, distress is a unique signal, and children are highly sensitive to others’ wellbeing. At 9 months of age children prefer prosocial actions over harmful ones; by the end of their first year they begin to comfort others; and by their second year of life, they engage in helping behavior [13, 22, 23].

Thus, while social identity influences social motivation in children including increased loyalty, sharing, and positive attitudes towards ingroup members [21, 24], it is unclear whether empathic helping is similarly prone to ingroup bias at young ages. Furthermore, encouragingly, children are more malleable than adults in their biases towards outgroup members[25]. Several studies have found that in humans, exposure to a diverse environment during childhood is associated with reduced biases into adulthood [26-29]. For example, unlike infants raised by families of their own race, infants in a multi-racial community do not prefer same-race faces [30]. Ingroup vs. outgroup categorization is thus flexible during human development. Yet, critically, the neural basis of the development of prosocial biases remains undefined, and could provide key insights into the flexibility of this biological mechanism.

Animal models have proved useful in the study of neural circuits underlying prosocial behavior. During the helping behavior test (HBT), adult rats who are exposed to a distressed trapped rat are motivated by that distress [31] to open a restrainer and release the trapped rat, even in the lack of social contact [32], demonstrating empathic helping. However, this prosocial behavior is socially selective, as rats release others from their own genetic strain, but do not help rats from unfamiliar strains[33, 34]. Furthermore, 2 weeks of co-habitation with a member of an unfamiliar strain caused a pro-social shift towards strangers of that strain, indicating that for rats, group membership is flexibly determined by social experience [33, 34]. The HBT is thus a good model for studying the neural processes involved in social bias for empathic helping in rats. Indeed, we found that neural regions typically associated with empathy, as well as reward, were active in rats following the HBT with trapped ingroup members [33]. In contrast, rats tested with outgroup members only showed activity in empathy networks. This pattern was not observed for non-trapped others, or for a non-social reward. Thus, while rats typically activate regions associated with negative arousal during the HBT, activity in the reward & motivation system is selectively associated with the presence of ingroup members and is predictive of helping.

While we have studied the neural bases of prosocial biases in adult animals, they have not yet been explored within a developmental context. Here we turned to young rats to examine the way adolescent brains respond to ingroup and outgroup members in distress during the HBT. We found that adolescent rats consistently released trapped outgroup members, in stark contrast to adults. Distinct patterns of movement and social interactions for ingroup and outgroup members suggest adolescents distinguish between the two conditions. After a final HBT session, a neural activity marker, the immediate early gene c-Fos, was analyzed to identify the neural networks associated with prosocial behavior. Distinct patterns of neural activity associated with each condition were observed and may underlie the generalized helping observed in adolescents compared to adults. In general, adolescents activated similar regions as adults during the HBT, reinforcing the participation of empathy and reward regions in this task. However, the hippocampus of adolescents was less active than adults, while activity in the orbitofrontal cortex was elevated. These findings suggest that the response to distress in adults may be inhibited by activation of circuits that respond to social category information. Overall, our findings demonstrate that adolescent rats do not show a similar ingroup bias as adults and display altered activity in networks of social mapping and reward valuation

## Results

### Adolescent rats, unlike adults, do not demonstrate an ingroup bias for prosocial behavior

Rats were tested in the helping behavior test (HBT), a simple test where rats can learn to open the door to a restrainer and release a conspecific trapped inside, as previously described in [32]. For hourly daily sessions over a two-week period, rats were given the opportunity to open the restrainer; they were not trained beforehand or rewarded in any way other than the reward afforded by door-opening and any subsequent social interaction. Here, helping behavior was studied in adolescent Sprague-Dawley (SD) rats (p32 days old) tested with age-matched SD cagemates (‘adolescent ingroup’, n=13), or with age-matched rats of the unfamiliar black-caped Long-Evans (LE) strain (‘adolescent outgroup’ n=8 Fig. 1A-B). Adolescent helping was compared to adult rats (p60-p90) tested with the same protocol (‘adult ingroup’, n=8 & ‘adult outgroup’, n=16). Part of this dataset was previously published in [33].

**Fig. 1.**
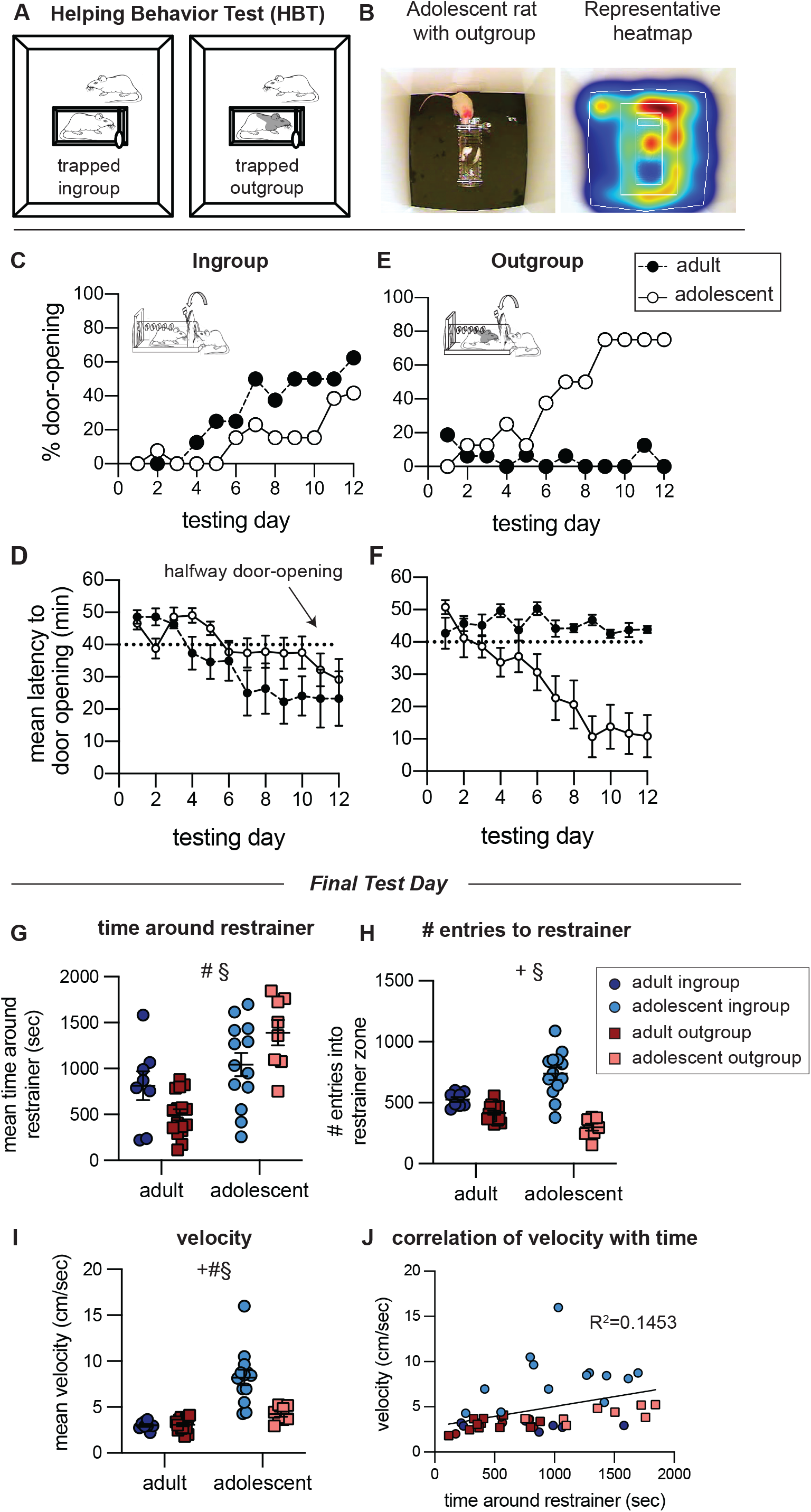
(above). Helping behavior for adult and adolescent rats. Adolescent rats, unlike adults, do not demonstrate an ingroup bias for prosocial behavior. (A) Diagram of the helping behavior test. (B) Representative movement pattern of an adolescent tested with an outgroup member depicted by a heatmap of the rat’s location along the session. (C-F) Helping behavior is expressed by % of door-openings and latency to open for the ingroup (C-D) and outgroup (E-F) across testing sessions. The dashed line indicates the half-way door-opening by the experimenter. (G-J) Analysis of movement patterns in the final testing session, including: (G) The time rats spent near the trapped rat, (H) the number of entries into the zone around the restrainer and (I) average velocity. (J) Time around restrainer was correlated with activity levels. 2-way ANOVA: + main effect of group identity, # main effect of age, § significant interaction between age and group identity.

Like adults, adolescent rats tested with ingroup members were motivated to release their trapped cagemates, as expressed by a significant increase in percent door-openings (Cochrans’ Q, p<0.01) and reduced latency to door-opening (Friedman, p<0.05) along the days of testing (Fig. 1C-D, Movie S1). Strikingly, unlike adults, adolescent rats robustly released trapped outgroup members, as expressed by a significant increase in the percent of door-openings (Cochrans’ Q, p<0.001) and decreased latency to open the restrainer door (Friedman, p<0.01, Fig. 1E-F, Movie S2). Nearly all rats in this condition (n=6/8) consistently released the trapped outgroup member, as opposed to 0/16 in the adult condition. The percent of door-openings did not increase in the adult outgroup condition, and door-opening behavior was rarely observed (Cochrans’ Q, Friedman, p>0.05). This unexpected finding demonstrates that the lack of prosocial motivation towards outgroup members emerges after early adolescence or in adulthood.

### Adolescent rats interact differently with ingroup and outgroup members

An unexpected finding was that adolescent rats were less successful at helping trapped ingroup members compared to adults. Only 4/13 adolescent rats became consistent openers by the end of testing, compared to 6/8 adult rats. This could point to reduced motivation to release trapped cagemates. However, both movement data and increased neural activity (described later) suggest they were highly motivated to do so. On the final testing day the restrainer was latched so that all rats had an objectively similar experience of being in the presence of a trapped conspecific for the entire session length. On this final test day, adolescents in the ingroup condition spent a similar amount of time around the trapped rat as the adolescent outgroup rats yet they entered the zone around the restrainer more frequently and were more active than the outgroup condition (ANOVA, p<0.01, Bonferroni p<0.01, Fig. 1G-I, Movie S3). Thus, despite lower rates of door-opening for adolescent ingroup than outgroup members, adolescents in the ingroup condition demonstrated movement patterns reflective of high motivation to release the trapped cagemate. In general, as is typically observed, adolescents in both conditions were more active than adults (Fig 1I). They also spent more time near the trapped rat than did adults on the final session (ANOVA, p<0.01, Bonferroni p<0.01, Fig. 1G-H), suggesting that a social stimulus is more salient for adolescents. Across all groups, activity was directed at the trapped rat; there was a positive correlation between activity and time near the restrainer (Pearson’s, p<0.01, Fig 1J), and rats were observed circling the restrainer as demonstrated in Fig. 1B and Movie S3. Combined, these data suggest that adolescents tested with cagemates were motivated, but less successful at learning the door-opening task than the adolescents tested with outgroup members. Future studies will be needed to explore the possible processes involved in this finding.

To further explore the motivational state of adolescents with trapped ingroup and outgroup members, social interactions immediately after door-opening were quantified on the day before the last session (the final day where social interaction was afforded, Fig. 2A, see methods). In line with the movement data, adolescents interacted with the freed conspecific more than adults (ANOVA, main effect of age, p<0.05), reinforcing the increased salience of social interaction for adolescents. Adolescents in the outgroup condition also showed the greatest number of interactions (Bonferroni, p<0.001, Fig. 2B). Yet the type of interaction was markedly different for adolescent ingroup and outgroup pairs: playfighting emerged as the predominant interaction in the adolescent ingroup condition (Bonferroni p<0.001, Fig. 2C), whereas non-play interactions, including anogenital sniffs, were significantly higher in the adolescent outgroup condition (Bonferroni p<0.001, Fig. 2D). Aggressive behaviors such as biting were rarely seen in any group and did not differ across the adolescent conditions (Fig. 2E). Thus, even on the final days of testing, rats behaved differently with ingroup and outgroup members, indicating they could distinguish between these social identities.

**Fig. 2.**
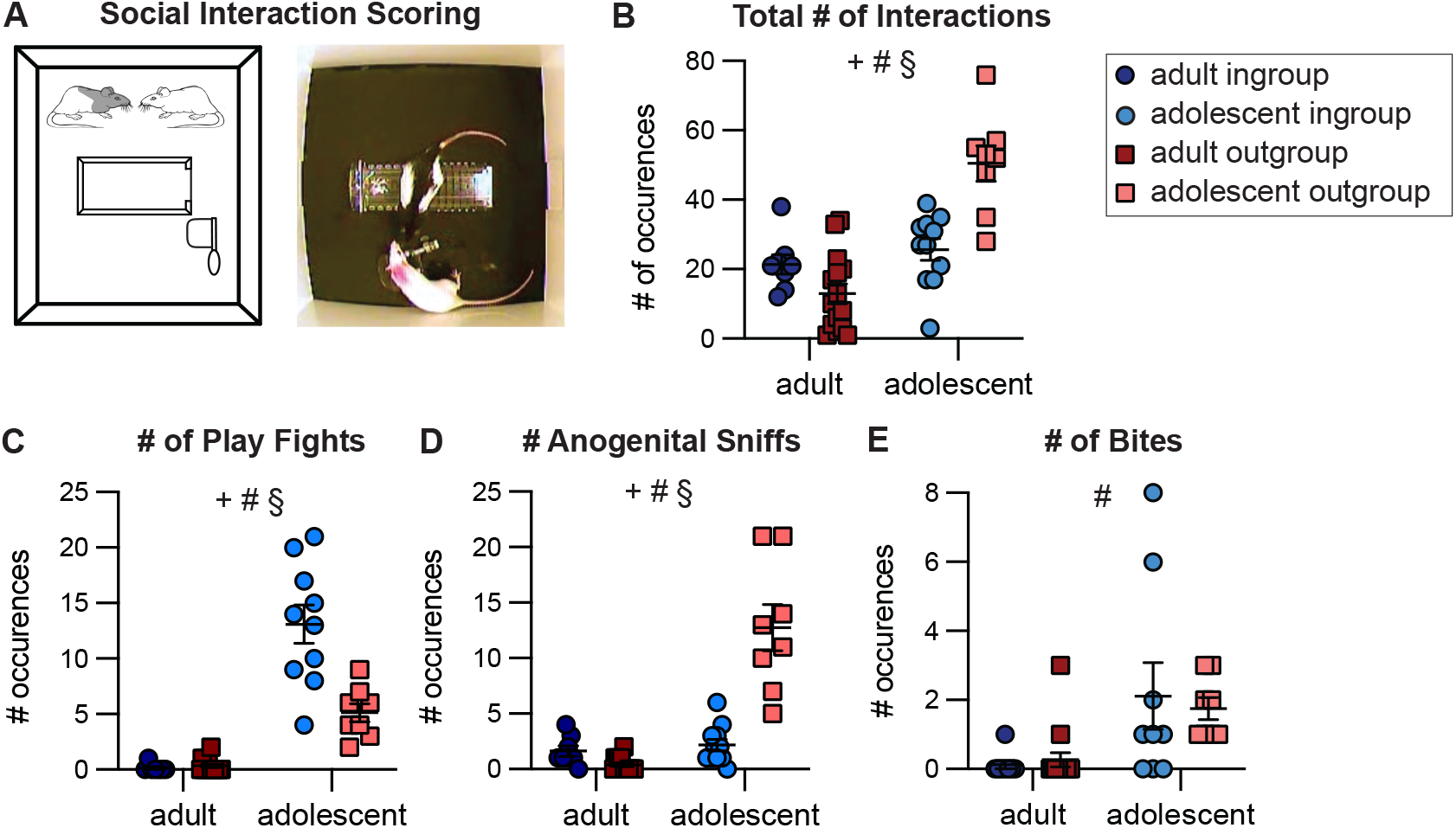
Adolescent rats display different types of social interaction depending on group identity. (A) Diagram and representative image of social interaction between an adolescent SD and LE rat. (B) Compared to adults, adolescents had a higher number of total social interactions scored within the 5-minute period. This includes all types of interactions, including play fighting, touching and investigations. (C) The number of play fights was highest in adolescents tested with cagemates. (D) The number of investigative anogenital sniffs was highest in adolescents tested with strangers. (E) The number of bites did not differ between adolescent groups. 2-way ANOVA: + main effect of group identity, # main effect of age, § significant interaction between age and group identity.

Altogether, we take these data to indicate that adolescents were more motivated than adults to release and interact with the trapped rat. The differing behaviors between the adolescent ingroup and outgroup conditions suggest that the free rats were sensitive to the group identity of the trapped rat and may point to two different motivational states in these conditions, such as empathy vs. curiosity or a desire for social interaction. Importantly, even if rats of all ages experience less emotional contagion with outgroup members, adolescents, in contrast with adults, release the trapped rat, demonstrating prosocial motivation and lack of social bias.

### Neural activity patterns in the helping behavior test correspond with age and group membership

In order to map brain-wide activation associated with the HBT across development, the immediate early-gene c-Fos was quantified as an index of neural activity. c-Fos was measured immediately following the final testing session during which the restrainer was latched shut, reflecting neural activity of rats in the presence of a trapped ingroup or outgroup member (n=84 sampled brain regions per rat, Fig. 3A-D, see detailed methods in: [33].

**Fig. 3.**
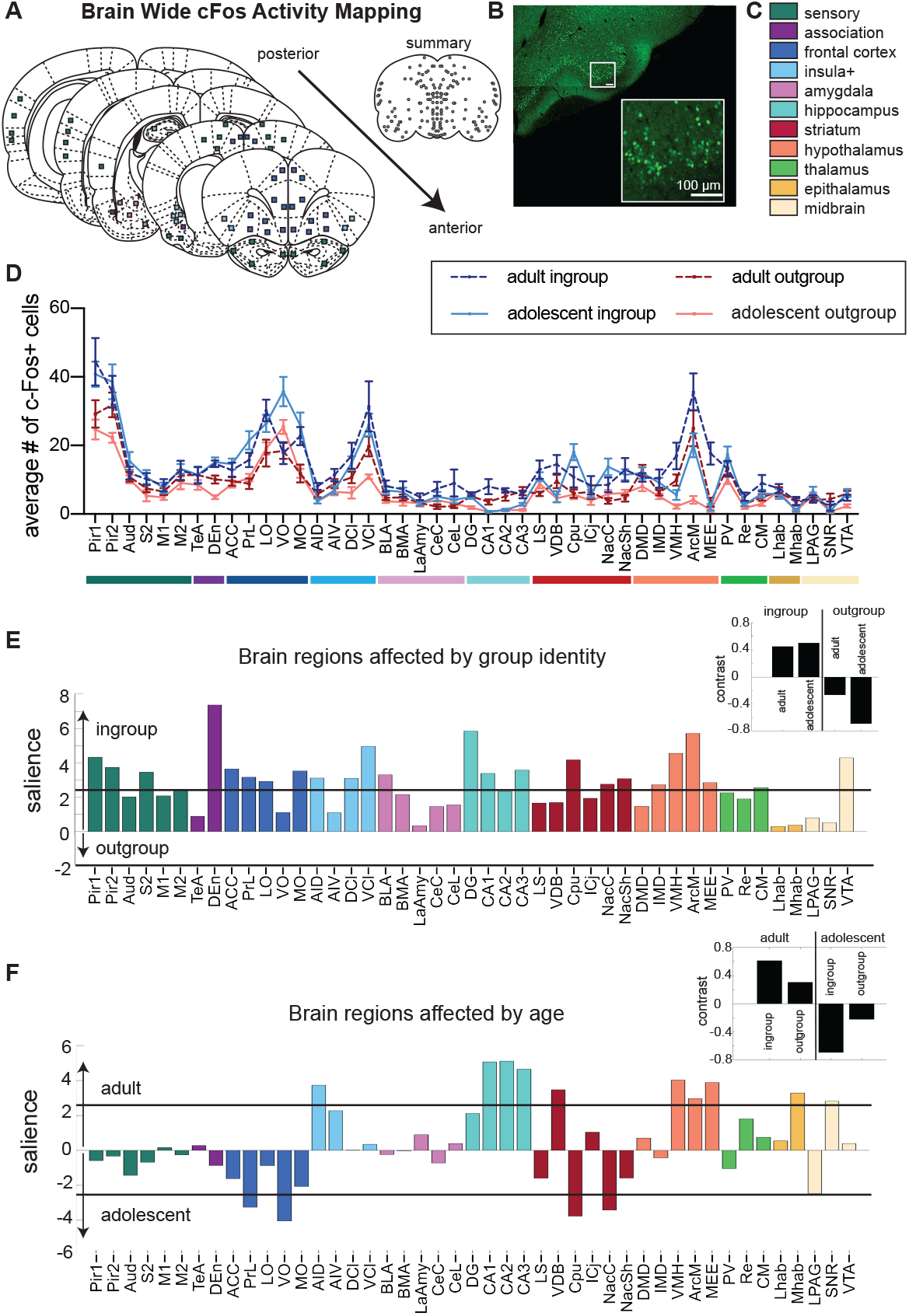
Neural activity associated with the helping behavior test. The brain-wide pattern of neural activity was determined by age and group identity. (A) Diagram of brain regions sampled for c-Fos expression. (B) A representative image of c-Fos signal sampled in the piriform cortex. (C) Legend of brain region categories coded by color. (D) Number of c-Fos+ cells per region (mean±SEM). Significant latent variables reveal that group identity (E) and age (F) determine neural activity patterns. The salience represents the z-score of boot-strapping tests, with regions crossing the black threshold lines significantly (p< 0.01) contributing to the contrast depicted in the inset (black bars). The directionality of the bars is congruent with the contrast graphs, as demonstrated by the arrows along the y-axis. All regions were more active for ingroup than outgroup members, but several regions (e.g. VO) were more active for adolescent than adult rats.

Two overarching patterns of neural activity were identified for the four HBT conditions using multivariate task partial least-square (PLS) analysis as previously described [33, 35, 36]. This analysis aims to identify patterns associated with each condition by maximizing the contrast between the tasks in a non-biased way. Two significant latent variables (LVs) emerged from data based on these four conditions, each one associated with a different pattern of neural activity, identified by permutation bootstrapping tests. One LV was associated with group identity (ingroup vs. outgroup, LV1, p<0.001, Fig. 3E), and the other was associated with age (adolescent vs. adult, LV2, p<0.001, Fig. 3F).

For both adolescent and adult rats, a distinct pattern of c-Fos activity emerged that was dependent on group identity. Specifically, exposure to a trapped ingroup member led to increased neural activity in a large number of brain regions, including in key regions previously observed to be uniquely active for ingroup relative to outgroup members in adults such as the nucleus accumbens (Nac), lateral septum (LS), prelimbic cortex (PrL), and medial orbitofrontal cortex (MO)[33].

Thus, regardless of age, the presence of trapped ingroup members recruits broad neural activity, indicating this is a more salient stimulus than a trapped outgroup member.

Whereas the first LV suggests most neural activity can be explained by group identity, the second LV emphasized overarching effects of age on neural activity, regardless of group identity. This LV can thus point to brain regions that are affected by development rather than social context; it revealed that adolescents displayed significantly reduced activity in the hippocampus, hypothalamus and dorsal anterior insula, as well as increased frontal activity compared to adults. Effects in the striatum were mixed, with reduced activity in the vertical limb of the diagonal band of Broca (VDB) and increased activity in the caudate putamen (Cpu) and nucleus accumbens shell (NacSh) for adolescents compared to adults (Fig. 3F).

To gain a better understanding of the interactions between group identity and age for each brain region, two-way ANOVAs with Bonferroni-corrected post-hoc tests were used to compare cFos+ cell numbers across the four HBT conditions (Fig 4, Table S1). As expected from the LVs above, some regions showed group-identity effects, others showed age effects, and some regions were impacted by both. Based on the significant LVs, results are presented for group identity (Fig 4A-B) and age (Fig 4C-D) separately; a full display of scatterplots is available in Fig. S1.

**Fig. 4.**
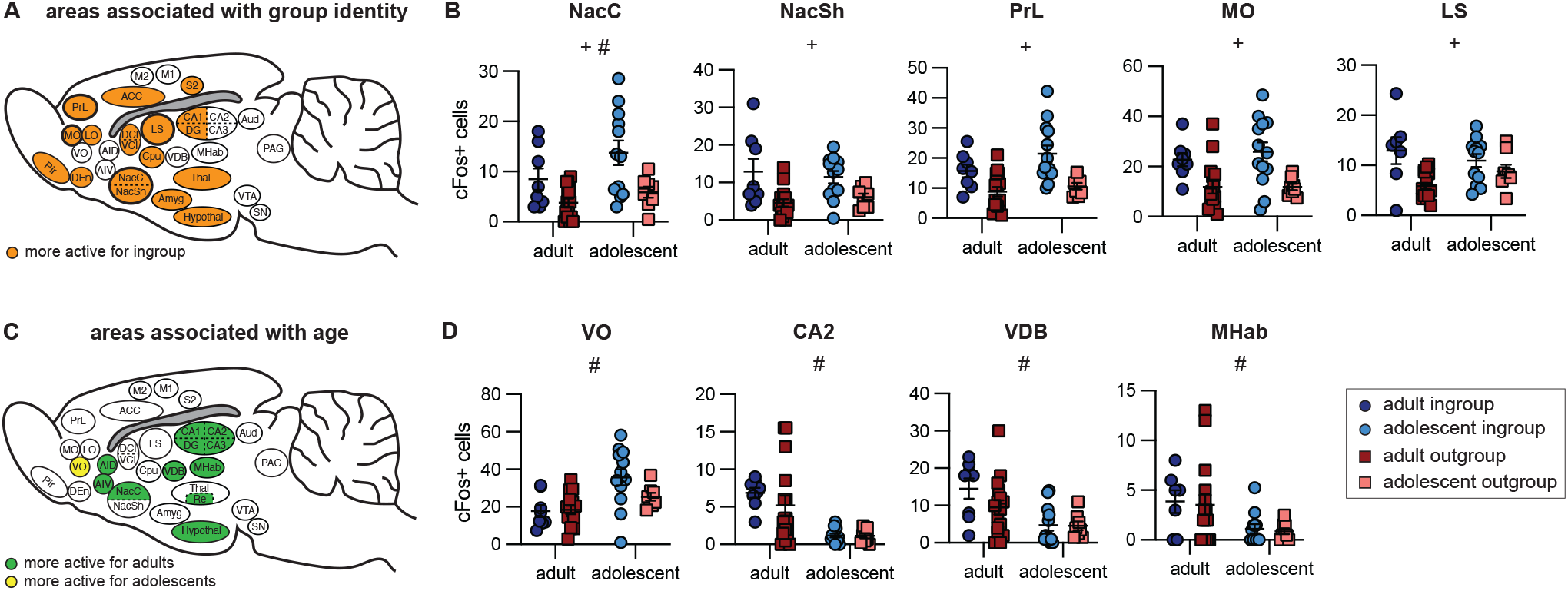
Neural activity of each condition, with main effects of group identity and age. Brain activity associated with group identity (A-B) and age (C-D) assessed by 2-way ANOVAs. (A) Brain diagram of all regions associated with group identity. All regions shown were more active for the ingroup than outgroup. (B) Scatterplots of five regions previously found to be uniquely active for adult ingroup compared to outgroup rats. Each region shows a main effect of group identity and adolescents display similar patterns as adults. (C) Brain diagram of all regions associated with age. Colored regions on the diagram represent areas more active for adults (green) or for adolescents (yellow). (D) Scatterplots of five of the seven brain regions that uniquely had a main effect of age but not group identity (not shown: AIV and CA3). All scatterplots can be found in Figure S1. 2-way ANOVA: + main effect of group identity, # main effect of age, § significant interaction between age and group identity.

First, we focused on regions of interest previously found to be more active for adult ingroup than outgroup members (based on: [33], (Fig 4A-B). These regions, the nucleus accumbens core (NacC), NacSh PrL, MO, and LS, all displayed main effects of condition (Table S1). Similar to adults, adolescents tested with ingroup members demonstrated increased c-Fos+ cell numbers in the nucleus accumbens core (NacC), PrL and MO (Bonferroni, p<0.05) relative to adolescents tested with outgroup members. In contrast, c-Fos numbers within the NacSh and LS were not significantly different across adolescent groups despite a main effect for group identity, pointing to developed sensitivity to group-identity in these regions. In addition, four of these five regions (all except the NacC) did not show a main effect of age, further indication that group identity rather than age drives these observed patterns of c-Fos activity. Conversely, to highlight age-associated effects, we examined regions contributing to the age LV but not the group LV in the PLS analysis, meaning these regions did not pass the significance threshold in the group identity salience plot (Fig 4C-D). We found that ventral orbitofrontal cortex (VO), medial habenula (MHab), VDB and CA2 of the hippocampus were more active for adults, whereas the VO was more active for adolescents (Fig. 4D). Thus, developmentally dependent increases in activity in these regions could indirectly explain the social selectivity in helping behavior observed in adults.

### Increased amygdala connectivity for adolescent non-openers

While adolescents in general were motivated to release the trapped rat, not all of them became successful helpers; these rats were classified as “non-openers” (see methods). When c-Fos levels were compared between openers and non-openers, a significant interaction emerged between opening and brain region (ANOVA, p<0.05), stemming from significantly more activity for non-openers in the ventral and lateral orbitofrontal cortex (OFC), piriform cortex, ventral claustrum (VCl) and medial arcuate hypothalamus (ArcM) (Bonferroni, p<0.05, Fig 5A). The increased activity in these regions for non-openers may stem from an increased motivation in the non-openers if, as posited above, adolescents in both groups were typically motivated to release the trapped rat. Further experiments will be needed to understand whether activity in these regions inhibits helping or reflects continued motivation.

**Fig. 5.**
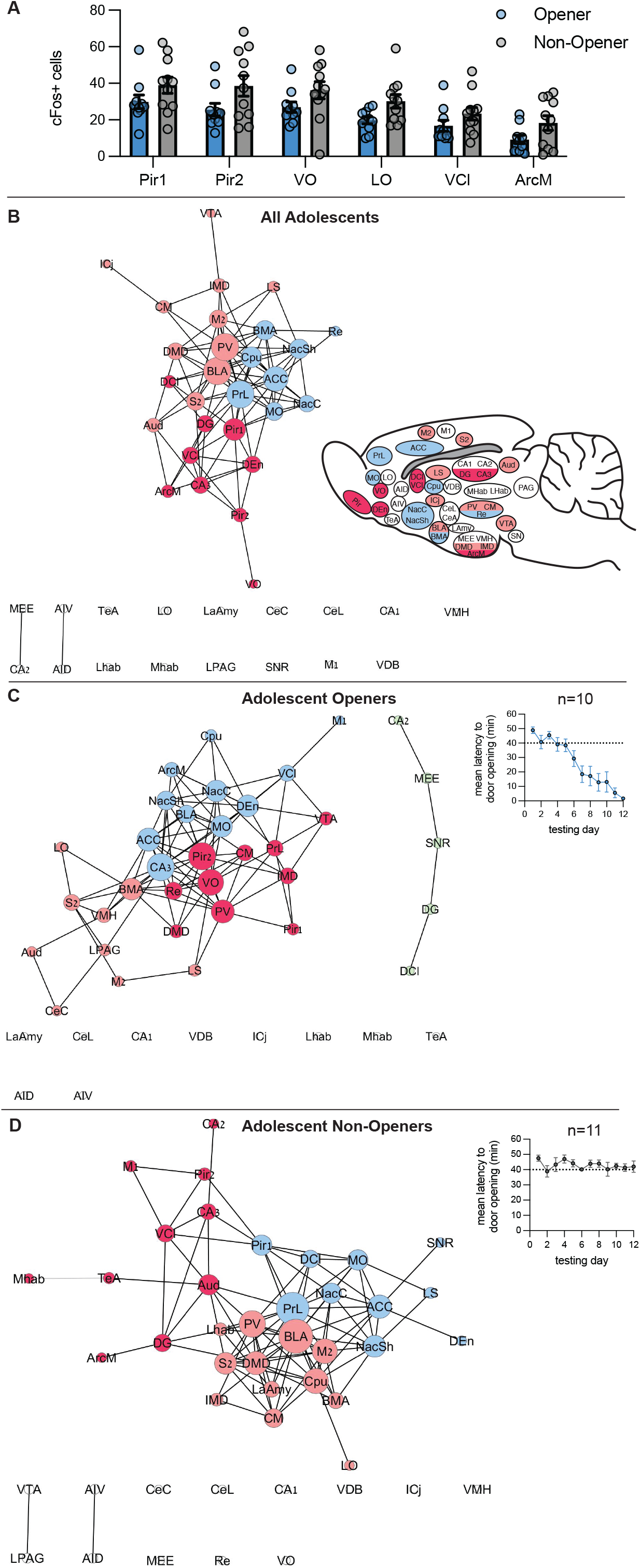
(above). Different neural patterns for opener and non-opener adolescent rats. (A) Brain regions with significantly higher levels of c-Fos for adolescent openers vs. non-openers are presented. (B-D) Network maps for adolescents tested in the HBT. (B) Network map for all adolescents, including rats in both the ingroup and outgroup condition. Inset: brain diagram colored by network clusters. (C) Network map for adolescent rats that became consistent openers. Inset: mean latency to door opening. (D) Network map for adolescent rats that did not consistently open across testing days. Inset: mean latency to door opening.

To gain insight into the way different adolescent brain regions interact during the HBT, network graph theory was used to generate functional connectivity maps based on c-Fos quantification. The networks present the top 10% correlated regions, based on a Pearson’s pair-wise correlation matrices (Fig. S2) and clustered using a Louvain algorithm, as previously reported in detail [33]. Note that this analysis highlights areas that are highly correlated with other brain regions; it does not describe overall activity levels. Using this method, a network map for all adolescent rats revealed 3 central clusters. Brain regions such as the PrL, MO and NAc, areas previously observed to be uniquely active in adult rats tested with cagemates, were also highly connected in one cluster of the network, alongside regions associated with empathy[4] such as the anterior cingulate cortex (ACC), suggesting that this network may be involved in the motivation to help in adolescents as well as in adults (Fig. 5B). Interestingly, mirroring the PLS and ANOVA findings, both the insula and the CA2 were not part of the adolescent network, and neither were areas associated with aversive responses (lateral and central amygdala, habenula, and others), indicating that these brain regions are not central to the adolescent response to a trapped cagemate.

We next examined the brain-wide patterns of functional connectivity by graphing the network maps for adolescent openers and non-openers. This analysis revealed that the main “motivational” cluster described above was largely conserved in both openers and non-opener networks, including connectivity between the MO, ACC and Nac (Fig. 5C-D). However, for non-openers a cluster containing amygdala regions emerged, including the basomedial, basolateral and lateral amygdala (BMA, BLA, LaAmy), and the habenula, indicating that connectivity in the amygdala may be detrimental to helping (Fig. 5D). Together, these findings demonstrate that common brain networks involved in reward and motivation were active in all adolescent rats, regardless of door opening behavior.

## Discussion

This study aimed to examine the neural development of social bias for prosocial behavior in adolescent rats. We found that in contrast with adults, adolescent rats did not show an ingroup bias, and instead helped trapped outgroup members, indicating that ingroup bias in rats emerges along development. One way to interpret the generalized helping in adolescent rats is a lack of sensitivity to group identity information due to later development of the neural circuits described above. However, the differences in movement patterns and social interactions provide behavioral evidence that rats do in fact distinguish between these social groups. An alternative explanation is that adolescents extend prosocial motivation to outgroup members, perhaps due to increased salience of social stimuli compared to adults, or a lack of threat arousal towards these adolescent outgroup conspecifics. In support of this explanation, we found increased exploratory interactions in the adolescent outgroup condition, compared to adults tested with outgroup members. As more affiliative interactions, such as playfighting, was observed for adolescent ingroup members, it is also possible that a different affective response was associated with each condition. While it is impossible to determine from these experiments if social reward, social investigation or empathic arousal was the main motivator for helping, the difference between adults and adolescents towards outgroup members is striking.

Adolescents tested in the HBT showed activation in a broadly dispersed neural network that responded preferentially to distressed ingroup members and was highly similar to that reported in adult rats[33, 37], as well as in humans[4]. This network includes regions in the sensory cortex, frontal cortex, ACC, anterior insula (AI), and reward and motivation areas like the claustrum, Cpu, Nac, hippocampus and hypothalamus. A different pattern of neural activity for adolescents relative to adult rats may indirectly explain the lack of social selectivity in prosocial motivation in adolescents. Adolescent rats showed decreased activation in several regions compared to adults. Specifically, CA2, VDB and MHab were significantly less active for adolescents and were not modulated by group identity. Furthermore, the LS, an area identified as more active for adults tested with ingroup than outgroup members, was similarly active for adolescent rats in both conditions. This suggests that the discrimination that occurs in the LS for group membership in adulthood is not apparent during adolescence. Interestingly, in newborn rat pups, specific layers of the LS have been shown to be active in response to the pup’s own mother and siblings, while other layers respond to another mother and her litter [38]. This suggests that at least some social identity information is represented in the LS in early life. It is possible then that the increased LS activity we see in adult ingroup vs. outgroup rats tested in the HBT represents a separate subpopulation that is specifically important for prosocial responding.

In general, sensitivity to social identity information has been observed in several brain regions including sensory cortex, dorsal medial PFC, LS, amygdala, and CA2 [39-44]. However, the source of social identity information as well as the directionality of information flow between these regions is unclear. Thus, selective responding based on social group could be represented in these regions due to downstream incorporation of social identity, which drives differential affective and motivational responses. Our results join with findings from other research groups and point to neural sensitivity to social information across multiple brain regions, including the hippocampus, amygdala, and striatum. [40, 45] The regions we and others have identified may be part of a neural circuit that connects information about social identity with motivated behavior. Both the VDB, a cholinergic basal forebrain region inhibiting magnocellular cells, and the LS are structurally connected to the hippocampus, and may be modulated by the CA2, a hippocampal region key to social mapping [46].

In particular, reduced hippocampal activation in adolescents may indicate a role for this region in the ingroup bias that emerges in adulthood. For instance, it is possible that social mapping is not distinctly defined in the adolescent brain. This idea is in line with research showing that discrimination based on social identity emerges in the amygdala in adulthood in humans [26] and mice [40]. Specifically, the CA2 is a possible target for future investigations. Indeed, in adolescents, the networks for differentiating social stimuli may not be well formed yet. Here, we examined functional networks in all adolescent rats and found that both the CA2 and insula regions were not functionally connected to the main network, reinforcing the finding that these regions are not centrally involved in the task before adulthood. Together, these findings support the hypothesis that CA2 becomes both more active and functionally connected to the rest of the brain in adulthood and participates in suppression of helping behavior towards non-affiliated others.

The OFC also emerged as an area of interest in this study. We previously found increased activity in the OFC for adult rats tested with both ingroup and outgroup members compared to baseline, with MO being significantly more active for the ingroup condition [33]. Here we found a similar trend, where MO and LO were significantly more active for adolescent ingroup members. Conversely, activity in the VO was not modulated by group identity, but it was impacted by age; the VO was the only region that was significantly more active for adolescents than adults. Interestingly, the VO was even more active in adolescent non-openers. As the OFC participates in processing rewards and evaluating outcomes[47], its specific modulation by group identity and success at helping may reflect involvement of the OFC in placing a value on the outcome of the trapped rat.

The current study faces several methodological limitations. First, there are limitations with using c-Fos staining; these have been extensively described in prior work [33]. Critically, c-Fos staining results in low temporal resolution, and thus, future work can expand upon the current study by using technology such as fiber photometry or activity targeted viral vectors to assess neural activity in adolescent rats undergoing the HBT. Higher temporal resolution will provide insight into neural activity during learning across the task, during door opening behavior and during subsequent social interactions, which we found differed according to group identity in adolescents. Here, our methodology using whole brain c-Fos adds to the growing validation of this type of unbiased approach in looking at brain activity in complex behaviors [48]. Our data suggest several key brain regions that may be responsible for helping behavior in adolescent rats. Future work will be able to expand on our findings to target specific regions and circuits, with the goal of artificially manipulating prosocial motivation across development. It is also important to note that the behavioral and neural findings here are from male rats. We are currently collecting data from both adult and adolescent female rats; how sex interacts with prosocial motivation will be critical to provide a more complete understanding of factors contributing to biases in helping behavior.

Our finding that adolescents help non-affiliated others opens up new areas for future investigation. Behaviorally, one hypothesis is that exposure to an outgroup member early in development may be sufficient to reduce biases in prosocial behavior. It will be worth exploring the bounds of this hypothesis; for example, is there a developmental window in which social context contributes to bias? Further, would a brief exposure of adolescent SD rats to LE strangers drive prosocial helping when tested as adults? Alternatively, adolescent rats may be driven to open for outgroup members due to social novelty or a desire for social investigation, as suggested from our social interaction data. Future studies will be able to directly address these hypotheses through manipulation of the early social environment and through manipulation of social interaction following door-opening. On a neural level, our findings suggest there may be a developmental trajectory of circuits that are not yet active in adolescents, including in the hippocampus and insula. Future work can test exactly when in development these brain regions become engaged in the larger network. In addition, future work could test the hypothesis that activation of hippocampal and/or insula regions are responsible for inhibition of helping outgroup members.

In conclusion, this study sheds light on the developmental basis of prosocial motivation and in-group bias. We demonstrate for the first time that adolescent rats are capable of helping behavior and help distressed others regardless of group identity. Further, we provide a window into the neural circuits associated with helping across development. Adolescent rats show a different pattern of neural activity during the HBT than adults; these differences may indirectly explain the lack of ingroup bias in adolescent rats. In particular, our results put a spotlight on the hippocampus and its role in group categorization, and suggest that in adults, CA2 activity may inhibit indiscriminate helping behavior. Overall, this study provides evidence for a developmental basis of prosocial helping across mammalian species and highlights a distinct neural response to the distress of affiliated others depending on age and group identity.

## General

We thank the following people for their help with this manuscript: Huanjie Sheng, Dominique Cajanding, Estherina Trachtenberg, Keren Ruzal and Einat Bigelman.

## Author contributions

Conceptualization IBB, JB, Investigation: IBB, JB, KL, JE, VG, NM, AZ, OP, KH, CP, PL, SC, CS, JC. Analysis: IBB, JB, HS, JE, VG, MB, AZ, OP, BK. Visualization & Writing: IBB, JB. Methodology: IBB. Resources & Funding: IBB, KL, DK. Supervision: IBB, DK.

## Data and materials availability

All data needed to evaluate the conclusions in the paper are present in the paper and/or the Supplementary Materials.

## STAR Methods

### Resource Availability

#### Lead Contact

Further information and requests for resources and reagents should be directed to and will be fulfilled by the lead contact, Inbal Ben-Ami Bartal.

#### Materials Availability

This study did not generate any new reagents or animal lines.

#### Data and Code Availability

All data have been uploaded in the following Open Science Framework depository (https://osf.io/6b2qc/) which is publicly available as of the date of this publication. DOIs are listed in the key resources table. This paper does not report original code. Any additional information required to reanalyze the data reported in this paper is available from the lead contact upon request.

### Experimental Model and Subject Details

#### Animals

Rat studies were performed in accordance with protocols approved by the Institutional Animal Care and Use Committee at the University of California, Berkeley. Rats were socially housed in cages of two same sex individuals, in a temperature (22-24C) and humidity controlled (55% relative humidity) animal facility, on a 12:12 light:dark cycle (lights on at 07:00). Food and Water was provided *ad libitum*. All testing was done in the rat’s light cycle. In total, 45 rats were tested across all experiments. For experiments with adults, male Sprague-Dawley rats (age postnatal day (p) 60-p90 days) were used as the free & trapped ingroup rats (Charles River, Portage, MI). Adult male Long-Evans rats were used as trapped outgroup rats (Envigo, CA). For experiments with adolescents, Sprague-Dawley (Charles River) rats were born in-house at UC Berkeley. Animals were separated by sex and weaned at p21, then were housed in pairs one week later at p28. Male Long-Evans rats (p28) housed in pairs were purchased from Charles River, as our Long-Evans breeders did not get pregnant as expected. All rats that were ordered were allowed a minimum of 5 days to acclimate to the facility prior to beginning testing. Trapped and free rats were of the same sex and age. Sprague Dawley animals were assigned to one of two experimental groups: they were either tested with cagemates (ingroup) or with Long-Evans strangers (outgroup).

### Method Details

#### Helping Behavior Test (HBT)

The helping behavior test (HBT) was performed as described previously [32]. Briefly, animals underwent five days of handling prior to starting the HBT. In addition to handling, on days 2-4, animals were given 30-minute habituation sessions where they were placed in an empty arena with their cagemate. On day 5, animals underwent a 15-minute open field task in the same arenas, one animal at a time. For the HBT, rats were tested in 60-minute sessions over a 12-day period.

On each day, rats were placed into arenas with either a trapped Sprague-Dawley rat (‘ingroup’) or Long-Evans rat (‘outgroup’) inside a restrainer located at the center of the arena. As in prior work, if the free rat did not open the restrainer after 40 minutes, the door was opened half-way by the experimenter. Both rats remained in the arena for the full hour. If the free rat opened the door before the half-way opening it was counted as a door-opening. After the initial 12 days, following a delay of typically one week, rats underwent three more test days. On the last day of testing, the restrainer was latched shut throughout the 60-minute session and rats were perfused within 30 minutes of completing behavioral testing. ‘Openers’ were defined as rats who opened the restrainer on at least two of the last three sessions (prior to the final day where the restrainers were latched shut). Sessions were video recorded with a CCD color camera (KT&C Co, Seoul, Korea) connected to a video card (Geovision, Irvine, CA) that linked to a PC. Movement data were analyzed using Ethovision video tracking software (Noldus Information Technology, Inc. Leesburg, VA). All adolescents began the first day of restrainer testing at approximately p32, while adults began the HBT between ages p60-p90.

#### Social Interaction Scoring

Five minutes of behavior was analyzed immediately upon release using BORIS software (see Key Resources Table). For rats that did not open the restrainer after 40 minutes, these interactions occurred in the final 20 minutes of the session once the trapped rat released himself. Two major categories of social behavior were scored: 1) play fighting interactions, including pinning and wrestling, and 2) non-play interactions, including nose to nose and nose to body touching and anogenital sniffs. Several videos could not be scored to do video encoding and export errors.

#### Immunohistochemistry

On the last day of testing, animals were sacrificed within 90 minutes from the beginning of the session, at the peak expression of the early immediate gene product c-Fos. Rats were transcardially perfused with 0.9% saline and freshly made 4% paraformaldehyde in phosphate buffered saline (PBS). Brains were then sunk in 30% sucrose as a cryoprotectant and frozen at - 80°C. They were later sliced at 40 μm and stained for c-Fos, as has been previously reported.[33] Sections were washed with 0.1M tris-buffered saline (TBS; 3×5’), incubated in 3% normal donkey serum (NDS) in 0.3% TritonX-100 in TBS (TxTBS), then transferred to rabbit anti-c-Fos antiserum (ABE457; Millipore, 1:1000; 1% NDS; 0.3% TxTBS) overnight. Sections were then washed in 0.1M TBS (3×5’), and incubated in Alexa Fluor 488-conjugated donkey anti-rabbit antiserum (AF488; Jackson, 1:500; 1% NDS; 0.3% TxTBS). Sections were then briefly washed in 0.1M TBS again (3×5’). Sections were further stained in DAPI (1:40,000), then washed for an additional 15 minutes (3×5’). Lastly, all slides were coverslipped with DABCO, dried overnight and stored at 4°C until imaged.

Immunostained tissue was imaged at 10x using a wide field fluorescence microscope (Zeiss AxioScan) and was processed in Zen software. Regions of interest (250 × 250μm squares) were placed across the whole brain, as described in[33]. A custom written script in ImageJ V2.0.0 (National Institute of Health, Bethesda, MD) was used to quantify immunoreactive nuclei, followed by manual checks and counting by multiple individuals who were blind to condition; consistency for counts across individuals was verified by a subset of samples. The threshold for detection of positive nuclei was set at a consistent level for each brain region, and only targets within the size range of 25–125 mm^2^ in area were counted as cells. Manual verification was targeted at identifying gross errors in the ImageJ scripts. For instance, in some cases the script falsely identified > 100 cells within the counting square; this usually occurred when there was high background staining. This type of error occurred in ∼15% of the samples, which were then manually corrected. 39 values for cell counts were removed from the dataset as outliers. Outliers were defined as those that were more than two standard deviations higher or lower than the group mean and further fell outside of the observed range for all conditions.

### Quantification and Statistical Analyses

Statistical details can be found within the Results section. In all written description and figures, n represents the number of animals in each condition. All means are reported as mean ± SEM. Statistical analyses described below were performed using MATLAB, SPSS, and Graphpad Prism.

#### Task Partial Least Square (PLS) analysis

Task PLS is a multivariate statistical technique that has been used to identify optimal patterns of activity that differentiate conditions [49, 50]. Task PLS is used in the analysis of brain region activity to describe the relationship between experimental conditions and correlated activity. PLS identifies similarities and differences between groups by locating regions where activation varies with the experimental condition. Through singular value decomposition, PLS produces a set of mutually orthogonal latent variable (LV) pairs. One element of the LV depicts the contrast, which reflects a commonality or difference between conditions. The other element of the LV, the brain region salience, identifies brain regions that show the activation profile across tasks, indicating which brain areas are maximally expressed in a particular LV.

Statistical assessment of PLS was performed by using permutation testing for latent variables (LVs) and bootstrap estimation of standard error for the brain region saliences. For the LV, significance was assessed by permutation testing: resampling without replacement by shuffling the test condition. Following each resampling, the PLS was recalculated. This was done 500 times in order to determine whether the effects represented in a given LV were significantly different than random noise. For brain region salience, reliability was assessed using bootstrap estimation of standard error. Bootstrap tests were performed by resampling 500 times with replacement, while keeping the subjects assigned to their conditions. This reflects the reliability of the contribution of that brain region to the LV. Brain regions with a bootstrap ratio greater than 2.55 (roughly corresponding to a confidence interval of 99%) were considered as reliably contributing to the pattern. Missing values were interpolated by the average for the test condition. An advantage to using this approach over univariate methods is that no corrections for multiple comparisons are necessary because the brain region saliences are calculated on all brain regions in a single mathematical step.

#### Network analysis

Network graphs were generated by first obtaining a correlation matrix of c-Fos activity between all brain regions (using pairwise Pearson correlation coefficients). The top 10% of correlations were presented in a graphic form. This cutoff threshold of 10% was determined based on scale-free network characteristics in prior work [33] and used here for comparability. Correlation values higher than the cutoff were set to one and the corresponding brain regions greater than 1 were considered connected to the network.

#### Other Statistical Tests

In addition to the PLS analysis described above, two-way ANOVAs were conducted on the c-Fos data to compare the four HBT conditions and to assess main effects of age (adult vs. adolescent) and group identity (ingroup vs outgroup). 2-way ANOVAs were also used to compare the pattern of animals’ movements during testing. Bonferroni post hoc corrections were used following all ANOVAs. Changes across days to helping behavior, including % door-opening and latency to door-opening, were examined using the non-parametric Cochran’s Q test and Friedman test respectively.

## Key resources table

| REAGENT or RESOURCE | SOURCE | IDENTIFIER |
| --- | --- | --- |
| **Antibodies** |  |  |
| **Rabbit anti-cFos primary antibody** | Millipore Sigma | Millipore: ABE457;<br>RRID: AB_2631318 |
| **Donkey anti-rabbit IgG Alexa Fluor 488 secondary antibody** | Jackson ImmunoResearch Labs | Cat#: 711-545-152;<br>RRID: AB_2313584 |
| **Deposited data** |  |  |
| https://osf.io/6b2qc/ |  |  |
| **Experimental models: Organisms/strains** |  |  |
| **Sprague-Dawley Rat** | Charles River Labs | Charles River 001;<br>RRID:<br>RGD_10395233 |
| **Long-Evans Rat** | Envigo | Envigo: HsdBlue:LE;<br>RRID: RGD_5508398 |
| **Software and algorithms** |  |  |
| **MATLAB** | Mathworks<br>(https://www.mathworks.com) | RRID: SCR_001622 |
| **SPSS** | IBM | RRID:SCR_019096 |
| **Zeiss ZEN 2 (Blue)** | Zeiss | RRID: SCR_013672 |
| **Fiji ImageJ** | NIH<br>(https://imagej.net/Fiji/Downloads); Schneider et al., 2012 | RRID: SCR_002285 |
| **Behavioral Observation Research Interactive Software Project (BORIS)** | https://edspace.american.edu/openbehavior/project/boris/ | RRID:SCR_021434 |
| **GraphPad Prism** | http://www.graphpad.com/ | RRID: SCR_002798 |

## Supplemental Figures & Tables

**Fig S1.**
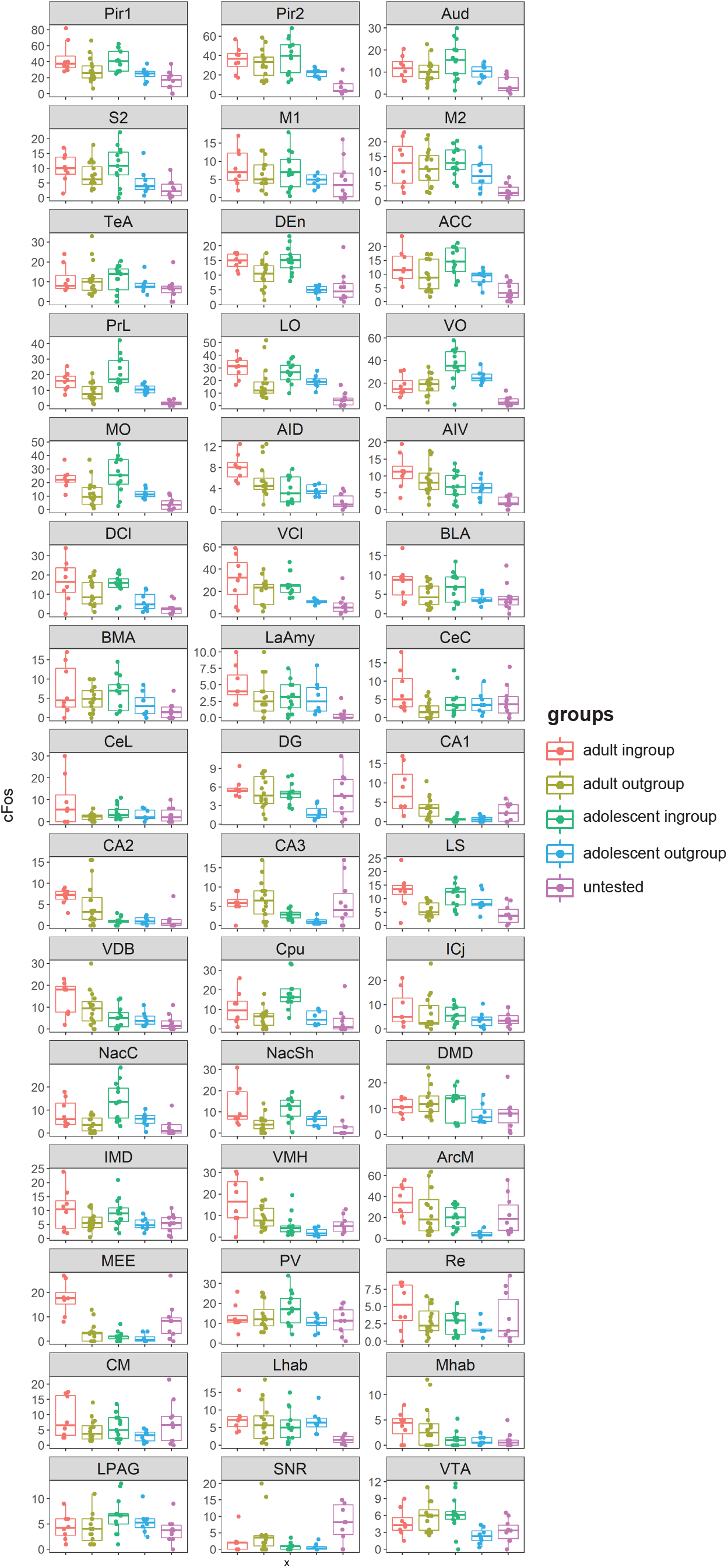
(above). Individual c-Fos expression for all regions and conditions. Box plots of c-Fos data in all brain regions across all test groups. Center bars mark the median. Lower and upper edges correspond to the 25th and 75th percentiles. Descriptions of the brain region abbreviations can be found in Table S1. Data points are jittered along the x-axis to avoid overlaps. X: experimental groups; Y: c-Fos^+^ cell numbers.

**Fig S2.**
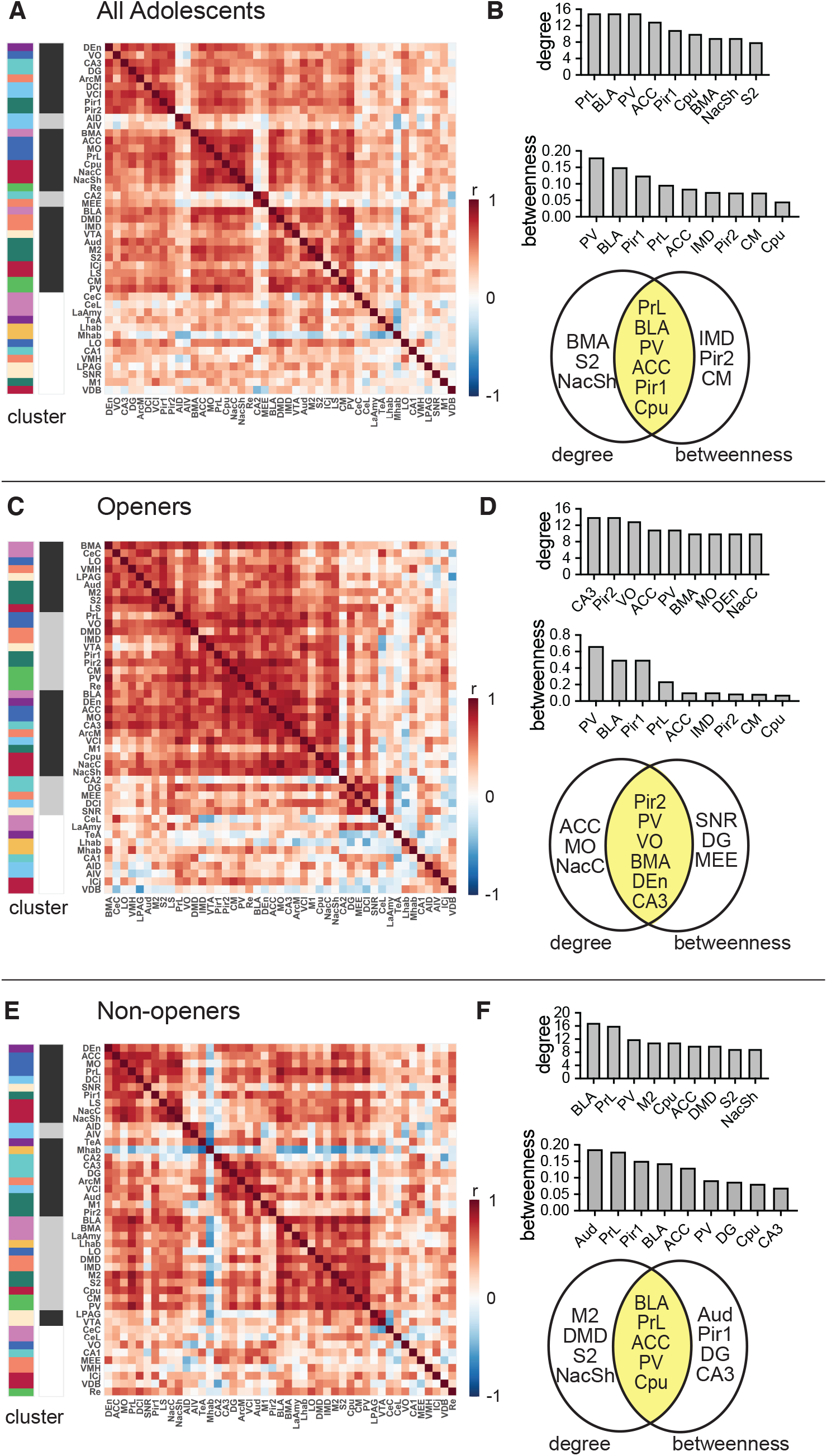
Correlation matrices. Pearson’s correlations across all brain regions in adolescent rats tested in the HBT. Correlation matrices (A,C,E) and central hubs (B,D,F) for (A-B): All adolescent rats, (C-D): Adolescent openers, and (E-F) Adolescent non-openers.

**Table S1.** 2-way ANOVA results. Main effects of group identity, age, and/or interaction between the two. The F statistic is shown, as well as statistically significant p-values in bold. \**p*<0.05, \*\**p*<0.01, \*\*\**p*<0.001, \*\*\*\**p*<0.0001.

| Brain Region | Group Identity | Age | Interaction |
| --- | --- | --- | --- |
| Pir1 | F (1, 40) = 11.63<br>**\*\* p=0.0015** | F (1, 40) = 0.8078 | F (1, 40) = 0.01069 |
| Pir2 | F (1, 41) = 1.946<br>**\* p=0.0303** | F (1, 41) = 0.5543 | F (1, 41) = 5.036 |
| Aud | F (1, 39) = 2.566 | F (1, 39) = 0.4879 | F (1, 39) = 1.104 |
| S2 | F (1, 39) = 5.648<br>**\* p=0.0225** | F (1, 39) = 0.2088 | F (1, 39) = 0.6472 |
| M1 | F (1, 38) = 2.608 | F (1, 38) = 0.7417 | F (1, 38) = 0.1015 |
| M2 | F (1, 42) = 2.593 | F (1, 42) = 0.3220 | F (1, 42) = 0.6461 |
| TeA | F (1, 38) = 0.4533 | F (1, 38) = 0.3985 | F (1, 38) = 0.5176 |
| DEn | F (1, 40) = 37.94<br>**\*\*\*\* p<0.0001** | F (1, 40) = 3.677 | F (1, 40) = 5.219<br>**\* p=0.0277** |
| ACC | F (1, 41) = 8.533<br>**\*\* p=0.0056** | F (1, 41) = 0.03591 | F (1, 41) = 0.8141 |
| PrL | F (1, 40) = 16.43<br>**\*\*\* p=0.0002** | F (1, 40) = 2.943 | F (1, 40) = 0.8072 |
| LO | F (1, 39) = 9.985<br>**\*\* p=0.0030** | F (1, 39) = 0.3394 | F (1, 39) = 0.2903 |
| VO | F (1, 40) = 2.031 | F (1, 40) = 13.20<br>**\*\*\*p=0.0008** | F (1, 40) = 2.549 |
| MO | F (1, 40) = 15.36<br>**\*\*\* p=0.0003** | F (1, 40) = 0.2487 | F (1, 40) = 0.2257 |
| AID | F (1, 40) = 1.922 | F (1, 40) = 13.55<br>**\*\*\* p=0.0007** | F (1, 40) = 1.732 |
| AIV | F (1, 41) = 1.470 | F (1, 41) = 5.303<br>**\* p=0.0264** | F (1, 41) = 0.7340 |
| DCI | F (1, 41) = 12.14<br>**\*\* p=0.0012** | F (1, 41) = 1.939 | F (1, 41) = 0.4127 |
| VCI | F (1, 41) = 12.52<br>**\*\* p=0.0010** | F (1, 41) = 3.991 | F (1, 41) = 0.2145 |
| BLA | F (1, 41) = 8.897<br>**\*\* p=0.0048** | F (1, 41) = 0.8785 | F (1, 41) = 0.03535 |
| BMA | F (1, 39) = 3.372 | F (1, 39) = 0.5021 | F (1, 39) = 0.006811 |
| LaAmy | F (1, 40) = 0.7826 | F (1, 40) = 0.9492 | F (1, 40) = 1.604 |
| CeC | F (1, 40) = 6.181<br>* **p=0.0172** | F (1, 40) = 0.008356 | F (1, 40) = 3.237 |
| CeL | F (1, 39) = 5.056<br>* **p=0.0303** | F (1, 39) = 1.232 | F (1, 39) = 3.135 |
| DG | F (1, 40) = 8.677<br>** **p=0.0053** | F (1, 40) = 10.97<br>** **p=0.0020** | F (1, 40) = 3.633 |
| CA1 | F (1, 39) = 4.987<br>* **p=0.0314** | F (1, 39) = 25.58<br>**** **p<0.0001** | F (1, 39) = 4.846<br>* **p=0.0337** |
| CA2 | F (1, 40) = 0.6554<br>ns | F (1, 40) = 20.93<br>**** **p<0.0001** | F (1, 40) = 0.6074<br>ns |
| CA3 | F (1, 38) = 0.09875 | F (1, 38) = 14.97<br>*** **p=0.0004** | F (1, 38) = 1.473 |
| LS | F (1, 39) = 11.24<br>** **p=0.0018** | F (1, 39) = 0.1116<br>ns | F (1, 39) = 3.227<br>ns |
| VDB | F (1, 40) = 1.589 | F (1, 40) = 13.02<br>*** **p=0.0008** | F (1, 40) = 1.414 |
| Cpu | F (1, 40) = 17.59<br>*** **p=0.0001** | F (1, 40) = 2.950 | F (1, 40) = 3.499 |
| ICj | F (1, 39) = 1.060 | F (1, 39) = 1.727 | F (1, 39) = 1.178e-006 |
| NAcC | F (1, 40) = 12.52<br>** **p=0.0010** | F (1, 40) = 4.243<br>* **p=0.0460** | F (1, 40) = 0.8062<br>ns |
| NAcSh | F (1, 40) = 14.73<br>*** **p=0.0004** | F (1, 40) = 0.001683<br>ns | F (1, 40) = 0.7023<br>ns |
| DMD | F (1, 40) = 0.8251 | F (1, 40) = 1.978 | F (1, 40) = 1.890 |
| IMD | F (1, 41) = 7.645<br>** **p=0.0085** | F (1, 41) = 0.5442 | F (1, 41) = 0.002939 |
| VMH | F (1, 41) = 6.212<br>* **p=0.0168** | F (1, 41) = 20.74<br>**** **p<0.0001** | F (1, 41) = 0.7902 |
| ArcM | F (1, 39) = 7.110<br>* **p=0.0111** | F (1, 39) = 13.41<br>*** **p=0.0007** | F (1, 39) = 0.2829 |
| MEE | F (1, 31) = 18.03<br>*** **p=0.0002** | F (1, 31) = 31.75<br>**** **p<0.0001** | F (1, 31) = 14.40<br>*** **p=0.0006** |
| PV | F (1, 39) = 2.405 | F (1, 39) = 0.03630 | F (1, 39) = 2.802 |
| Re | F (1, 40) = 5.066<br>**\* p=0.0300** | F (1, 40) = 6.502<br>**\* p=0.0147** | F (1, 40) = 0.7999 |
| CM | F (1, 41) = 7.399<br>**\*\* p=0.0095** | F (1, 41) = 3.557 | F (1, 41) = 0.1994 |
| Lhab | F (1, 39) = 0.002036 | F (1, 39) = 0.1812 | F (1, 39) = 0.4681 |
| Mhab | F (1, 38) = 0.08974 | F (1, 38) = 7.920<br>**\*\* p=0.0077** | F (1, 38) = 0.001485 |
| LPAG | F (1, 35) = 0.5708 | F (1, 35) = 2.982 | F (1, 35) = 0.1695 |
| SNR | F (1, 36) = 0.5972 | F (1, 36) = 3.947 | F (1, 36) = 0.8727 |
| VTA | F (1, 36) = 1.913 | F (1, 36) = 1.437 | F (1, 36) = 7.306<br>**\* p=0.0104** |

## References

1. van Schaik, C.P., and Burkart, J.M. (2011). Social learning and evolution: the cultural intelligence hypothesis. Philos Trans R Soc Lond B Biol Sci 366, 1008–1016.

2. Wilson, E.O. (2012). The Social Conquest of Earth, (New-York: Liveright Publishing Corporation).

3. Dugatkin, L.A. (1997). Cooperation among animals: an evolutionary perspective, (New York: Oxford University Press).

4. Decety, J., Bartal, I.B., Uzefovsky, F., and Knafo-Noam, A. (2016). Empathy as a driver of prosocial behaviour: highly conserved neurobehavioural mechanisms across species. Philos. Trans. R. Soc Lond B Biol. Sci 371.

5. Hamilton, W.D. (1964). The genetical evolution of social behaviour. I. J Theor. Biol 7, 1–16.

6. Hackel, L.M., Zaki, J., and Van Bavel, J.J. (2017). Social identity shapes social valuation: evidence from prosocial behavior and vicarious reward. Soc Cogn Affect Neurosci 12, 1219–1228.

7. Bernhard, H., Fischbacher, U., and Fehr, E. (2006). Parochial altruism in humans. Nature 442, 912–915.

8. Kavaliers, M., and Choleris, E. (2017). Out-Group Threat Responses, In-Group Bias, and Nonapeptide Involvement Are Conserved across Vertebrates: (A Comment on Bruintjes et al., “Out-Group Threat Promotes Within-Group Affiliation in a Cooperative Fish”). Am Nat 189, 453–458.

9. Fiske, S.T. (2002). What We Know Now About Bias and Intergroup Conflict, the Problem of the Century. Current directions in psychological science : a journal of the American Psychological Society 11, 123–128.

10. Andrews, J.L., Ahmed, S.P., and Blakemore, S.J. (2021). Navigating the Social Environment in Adolescence: The Role of Social Brain Development. Biol Psychiatry 89, 109–118.

11. Kilford, E.J., Garrett, E., and Blakemore, S.J. (2016). The development of social cognition in adolescence: An integrated perspective. Neurosci Biobehav Rev 70, 106–120.

12. Fuhrmann, D., Knoll, L.J., and Blakemore, S.J. (2015). Adolescence as a Sensitive Period of Brain Development. Trends Cogn Sci 19, 558–566.

13. Decety, J., Steinbeis, N., and Cowell, J.M. (2021). The neurodevelopment of social preferences in early childhood. Curr Opin Neurobiol 68, 23–28.

14. Charlesworth, T.E.S.B. M.R., (2021). The development of social group cognition: what infants and young children can teach us., 2nd Edition Edition, (Oxford Handbook of Social Congition).

15. Xiao, N.G., Wu, R., Quinn, P.C., Liu, S., Tummeltshammer, K.S., Kirkham, N.Z., Ge, L., Pascalis, O., and Lee, K. (2018). Infants Rely More on Gaze Cues From Own-Race Than Other-Race Adults for Learning Under Uncertainty. Child Dev 89, e229–e244.

16. Kinzler, K.D., Dupoux, E., and Spelke, E.S. (2007). The native language of social cognition. Proc Natl Acad Sci U S A 104, 12577–12580.

17. Shutts, K., Banaji, M.R., and Spelke, E.S. (2010). Social categories guide young children’s preferences for novel objects. Dev Sci 13, 599–610.

18. Shutts, K., Kinzler, K.D., McKee, C.B., and Spelke, E.S. (2009). Social information guides infants’ selection of foods. J Cogn Dev 10, 1–17.

19. Aboud, F.E. (2003). The formation of in-group favoritism and out-group prejudice in young children: are they distinct attitudes? Dev Psychol 39, 48–60.

20. Hailey, S.E.O. K.R., (2013). A social psychologist’s guide to the development of racial attitudes. Social and Personality Psychology Compass 7, 457–469.

21. Bigler, R.S., Jones, L.C., and Lobliner, D.B. (1997). Social categorization and the formation of intergroup attitudes in children. Child Dev 68, 530–543.

22. Gross, R.L., Drummond, J., Satlof-Bedrick, E., Waugh, W.E., Svetlova, M., and Brownell, C.A. (2015). Individual differences in toddlers’ social understanding and prosocial behavior: disposition or socialization? Front Psychol 6, 600.

23. Svetlova, M., Nichols, S.R., and Brownell, C.A. (2010). Toddlers’ prosocial behavior: from instrumental to empathic to altruistic helping. Child Dev 81, 1814–1827.

24. Over, H. (2018). The influence of group membership on young children’s prosocial behaviour. Curr Opin Psychol 20, 17–20.

25. Nelson, E.E., Jarcho, J.M., and Guyer, A.E. (2016). Social re-orientation and brain development: An expanded and updated view. Dev Cogn Neurosci 17, 118–127.

26. Telzer, E.H., Humphreys, K.L., Shapiro, M., and Tottenham, N. (2013). Amygdala sensitivity to race is not present in childhood but emerges over adolescence. J. Cogn Neurosci 25, 234–244.

27. X., Z., and S., H. (2013). Cultural experiences reduce racial bias in neural responses to others’ suffering. Cult. Brain 1, 34–46.

28. Heron-Delaney, M., Anzures, G., Herbert, J.S., Quinn, P.C., Slater, A.M., Tanaka, J.W., Lee, K., and Pascalis, O. (2011). Perceptual training prevents the emergence of the other race effect during infancy. PLoS One 6, e19858.

29. Anzures, G., Wheeler, A., Quinn, P.C., Pascalis, O., Slater, A.M., Heron-Delaney, M., Tanaka, J.W., and Lee, K. (2012). Brief daily exposures to Asian females reverses perceptual narrowing for Asian faces in Caucasian infants. J Exp Child Psychol 112, 484–495.

30. Bar-Haim, Y., Ziv, T., Lamy, D., and Hodes, R.M. (2006). Nature and nurture in own-race face processing. Psychol Sci 17, 159–163.

31. Ben-Ami, B. I, Shan, H., Molasky, N., Murray, T., Williams, J., Decety, J., and Mason, P. (2016). Pro-Social Behavior In Rats Requires An Affective Motivation. In BioRxiv.

32. Ben-Ami, B. I, Decety, J., and Mason, P. (2011). Empathy and pro-social behavior in rats. Science 334, 1427–1430.

33. Ben-Ami Bartal, I., Breton, J.M., Sheng, H., Long, K.L., Chen, S., Halliday, A., Kenney, J.W., Wheeler, A.L., Frankland, P., Shilyansky, C., et al. (2021). Neural correlates of ingroup bias for prosociality in rats. Elife 10.

34. Ben-Ami, B. I, Rodgers, D.A., Bernardez Sarria, M.S., Decety, J., and Mason, P. (2014). Pro-social behavior in rats is modulated by social experience. Elife 3, e01385.

35. Wheeler, A.L., Teixeira, C.M., Wang, A.H., Xiong, X., Kovacevic, N., Lerch, J.P., McIntosh, A.R., Parkinson, J., and Frankland, P.W. (2013). Identification of a functional connectome for long-term fear memory in mice. PLoS Comput Biol 9, e1002853.

36. Vetere, G., Kenney, J.W., Tran, L.M., Xia, F., Steadman, P.E., Parkinson, J., Josselyn, S.A., and Frankland, P.W. (2017). Chemogenetic Interrogation of a Brain-wide Fear Memory Network in Mice. Neuron 94, 363–374 e364.

37. Meyza, K.Z., Bartal, I.B., Monfils, M.H., Panksepp, J.B., and Knapska, E. (2017). The roots of empathy: Through the lens of rodent models. Neurosci Biobehav Rev 76, 216–234.

38. Clemens, A.M., Wang, H., and Brecht, M. (2020). The lateral septum mediates kinship behavior in the rat. Nat Commun 11, 3161.

39. Kohl, J., Babayan, B.M., Rubinstein, N.D., Autry, A.E., Marin-Rodriguez, B., Kapoor, V., Miyamishi, K., Zweifel, L.S., Luo, L., Uchida, N., et al. (2018). Functional circuit architecture underlying parental behaviour. Nature 556, 326–331.

40. Bergan, J.F., Ben-Shaul, Y., and Dulac, C. (2014). Sex-specific processing of social cues in the medial amygdala. Elife 3, e02743.

41. Raam, T., and Hong, W. (2021). Organization of neural circuits underlying social behavior: A consideration of the medial amygdala. Curr Opin Neurobiol 68, 124–136.

42. Kingsbury, L., Huang, S., Raam, T., Ye, L.S., Wei, D., Hu, R.K., Ye, L., and Hong, W. (2020). Cortical Representations of Conspecific Sex Shape Social Behavior. Neuron 107, 941–953 e947.

43. Baez-Mendoza, R., Mastrobattista, E.P., Wang, A.J., and Williams, Z.M. (2021). Social agent identity cells in the prefrontal cortex of interacting groups of primates. Science 374, eabb4149.

44. Carcea, I., Caraballo, N.L., Marlin, B.J., Ooyama, R., Riceberg, J.S., Mendoza Navarro, J.M., Opendak, M., Diaz, V.E., Schuster, L., Alvarado Torres, M.I., et al. (2021). Oxytocin neurons enable social transmission of maternal behaviour. Nature 596, 553–557.

45. Yao, S., Bergan, J., Lanjuin, A., and Dulac, C. (2017). Oxytocin signaling in the medial amygdala is required for sex discrimination of social cues. Elife 6.

46. Hitti, F.L., and Siegelbaum, S.A. (2014). The hippocampal CA2 region is essential for social memory. Nature 508, 88–92.

47. Simon, N.W., and Moghaddam, B. (2015). Neural processing of reward in adolescent rodents. Dev Cogn Neurosci 11, 145–154.

48. Turkheimer, F.E., Rosas, F.E., Dipasquale, O., Martins, D., Fagerholm, E.D., Expert, P., Vasa, F., Lord, L.D., and Leech, R. (2021). A Complex Systems Perspective on Neuroimaging Studies of Behavior and Its Disorders. Neuroscientist, 1073858421994784.

49. McIntosh, A.R., Bookstein, F.L., Haxby, J.V., and Grady, C.L. (1996). Spatial pattern analysis of functional brain images using partial least squares. Neuroimage 3, 143–157.

50. McIntosh, A.R. (1999). Mapping cognition to the brain through neural interactions. Memory 7, 523–548.

